# Gamma-aminobutyric acid treatment reduces estradiol-stimulated breast cancer cell growth and estradiol-induced breast tumorigenesis

**DOI:** 10.1101/2025.01.04.631336

**Authors:** Akihiro Nakamura, Sunao Tanaka, Haruna Shimizu, Hideki Hokazono, Masakazu Toi, Junji Itou

## Abstract

Gamma-aminobutyric acid (GABA) is a non-protein amino acid and contained in various foods, such as fruits and vegetables. Although various health benefits of GABA have been reported, GABA seems to have unidentified functions. In this study, we first show increase in intracellular GABA level in GABA-treated breast cancer cell lines, MCF-7 and T-47D, in amino acid analyses and immunofluorescence. In our experimental condition, GABA treatment reduced estradiol-stimulated cell growth in these cell lines. In addition, we performed mouse early breast cancer induction model system to investigate GABA function in mammary dysplasia formation. In the results, GABA administration reduced dysplasia formation, suggesting that GABA prevents breast tumorigenesis. This study demonstrates a novel function of GABA to maintain normal breast tissues.

## Introduction

Benefits of foods are not only for energy, source of body components and cultures, but also for health, for instance maintenance of physical ability and preventing diseases. Gamma-aminobutyric acid (GABA) is a non-protein amino acid, and was first found in the potato tuber as a substance which reacted to ninhydrin (1). Lactic acid bacteria produces GABA (2). GABA is contained in various fruits, vegetables, grains, honey and lactic acid-fermented foods (3, 4). In addition, GABA-enriched foods are offered (5).

In animals, various health benefits of GABA have been reported (2, 6, 7). Oral intake of GABA-containing fermented milk reduced systolic and diastolic blood pressures in patients with mild hypertensive, comparing to control patients who took acidified milk without GABA (8). GABA seems to protect liver tissues from injuries induced by D-galactosamine or alcohol (9, 10).

Breast cancer patients with higher GABA level than 89.3 μg/L in their tumors showed longer overall survival comparing to that with lower GABA level (11). However, the function of GABA is still controversial in the studies using cancer cell lines (12). GABA receptor signaling promotes metastasis in human and mouse breast cancer cells (13-15). On the other hand, GABA inhibits metastatic properties in triple-negative breast cancer cell lines, MDA-MB-468 (16).

Although GABA acts as an extracellular signaling molecule to activate GABA receptors, there are transporters which transport GABA molecules through membrane (17, 18), such as SLC6A6 and SLC6A8. Oral GABA intake increases plasma GABA level in healthy humans (19). In this study, we first determine whether GABA treatment increases intracellular GABA level in estrogen receptor positive breast cancer cell lines. In our condition, GABA inhibits cell growth stimulated by estradiol (E2), one of estrogen hormones. In addition, we show the preventive effect of GABA in E2-induced breast tumorigenesis in mouse early breast cancer induction model. Our study demonstrates a novel effect of GABA to maintain normal breast tissue.

## Results

### GABA treatment increases intracellular GABA level in breast cancer cell lines

To determine whether GABA treatment increases intracellular levels of GABA, we measured GABA levels with an amino acid analyzer in breast cancer cell lines, MCF-7 and T-47D. GABA signal was increased in cells incubated with PBS containing 1 μg/mL GABA for 0.5 h (Fig. 1AB). GABA signal was not observed in supernatant samples from cells treated with fixation solution whose GABA molecules were crosslinked and not moved to supernatants (Fig, 1B).

**Fig. 1.**
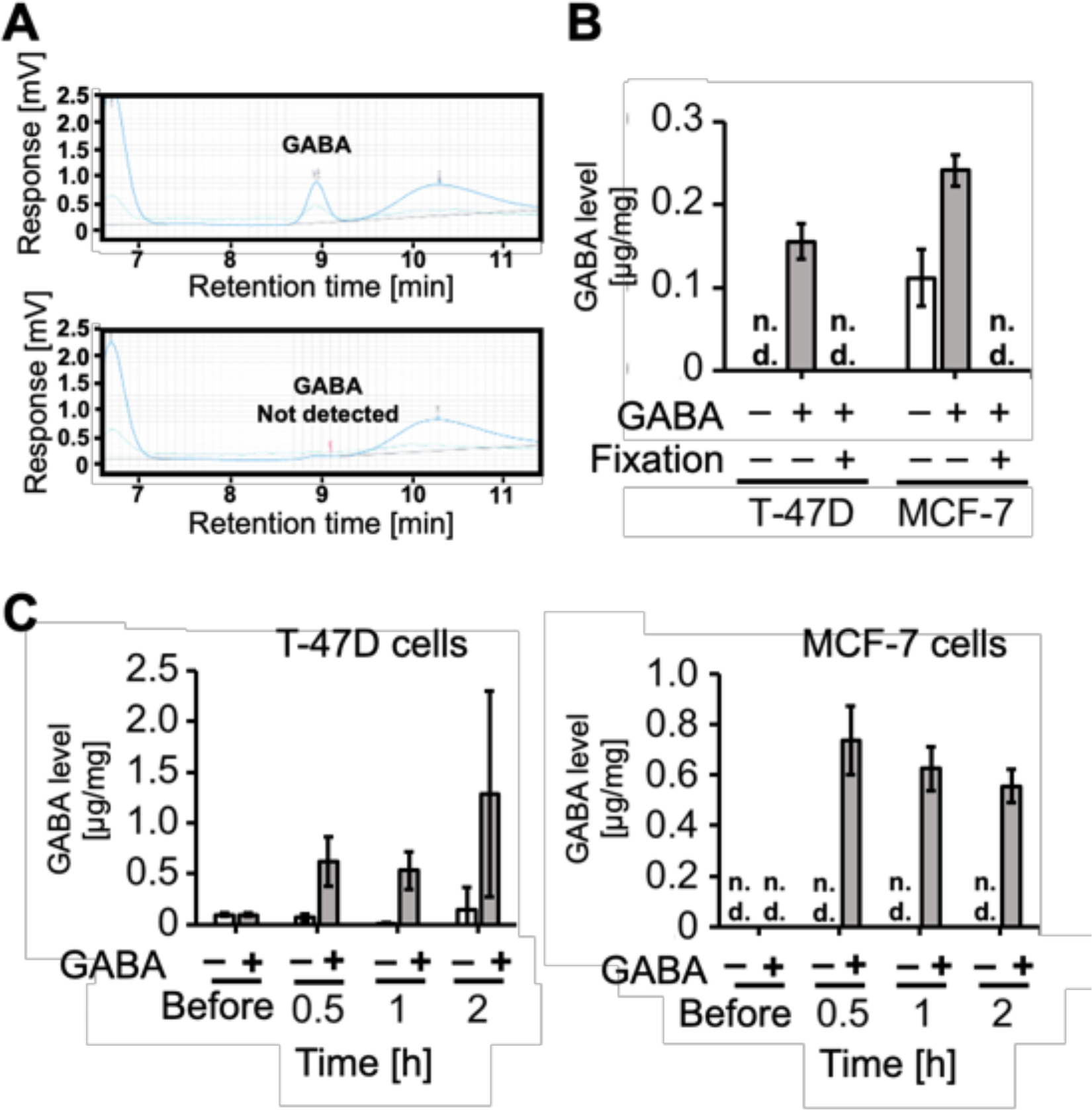
Detection of increased intracellular GABA levels by amino acid analyses. (A) Chromatograms of amino acid analyses are shown. T-47D cells were used. (B) Detected GABA levels are graphed. Cell fixation with formaldehyde was performed as negative controls. (C) The results of time course analyses are shown. Cells were incubated for 0.5, 1, 2 h with or without GABA. Values of before-treatment samples are shown as negative controls. n.d.: not detected.

To quantify intracellular GABA level in each sample, we measured the protein level of the trichloroacetic acid precipitated pellet of the same sample, and GABA signal was normalized by it. In the results, we observed increase in intracellular GABA levels in the cells incubated with GABA-containing PBS, comparing to cells incubated with PBS (Fig. 1C). These results indicate that GABA administration increases GABA level.

To visualize intracellular GABA, we performed immunostaining with a commercially available anti-GABA antibody. In confocal microscopy, we observed increase in GABA signal in GABA-treated cells (Fig. 2). Increased GABA signal was not detected in immunostainings without permeabilization in which anti-GABA antibodies did not penetrate into a cell and not react with intracellular GABA, indicating that our immunostaining with permeabilization detected increased intracellular GABA level in GABA treated cells. These results suggest that administered GABA molecules were incorporated into a cell.

**Fig. 2.**
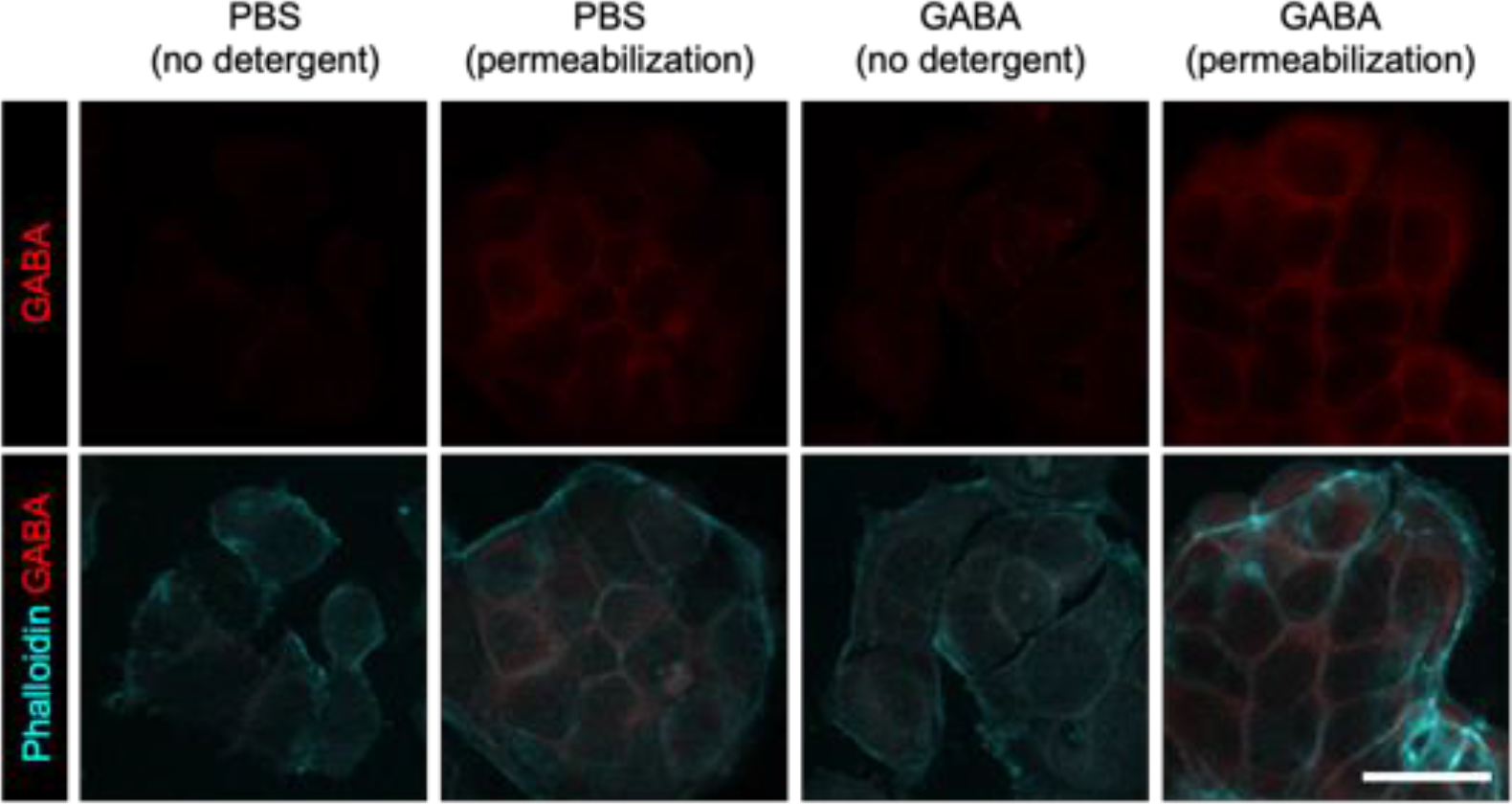
Increase in anti-GABA antibody immunoreaction in the inside of GABA treated cells. Confocal images of immunofluorescence with anti-GABA antibody are shown. MCF-7 cells were used. Cytoskeleton staining with phalloidin was performed to visualize the outlines of cells. Scale bar indicates 30 μm.

### GABA-treatment reduces cell growth in breast cancer cell lines

Although the function of GABA is largely unknown, GABA treatment reduces cell growth in cancers (20-22). To determine whether GABA treatment also reduces E2-stimulated cell growth of estrogen receptor-positive breast cancer cells, we used MCF-7 and T-47D cells (Fig. 3). In the results, E2 administration significantly increased cell numbers at day 3 and day 4, comparing to no E2 groups. GABA treatment reduced E2-stimulated cell growth, whereas not reduced in no E2 administered cell growth.

**Fig. 3.**
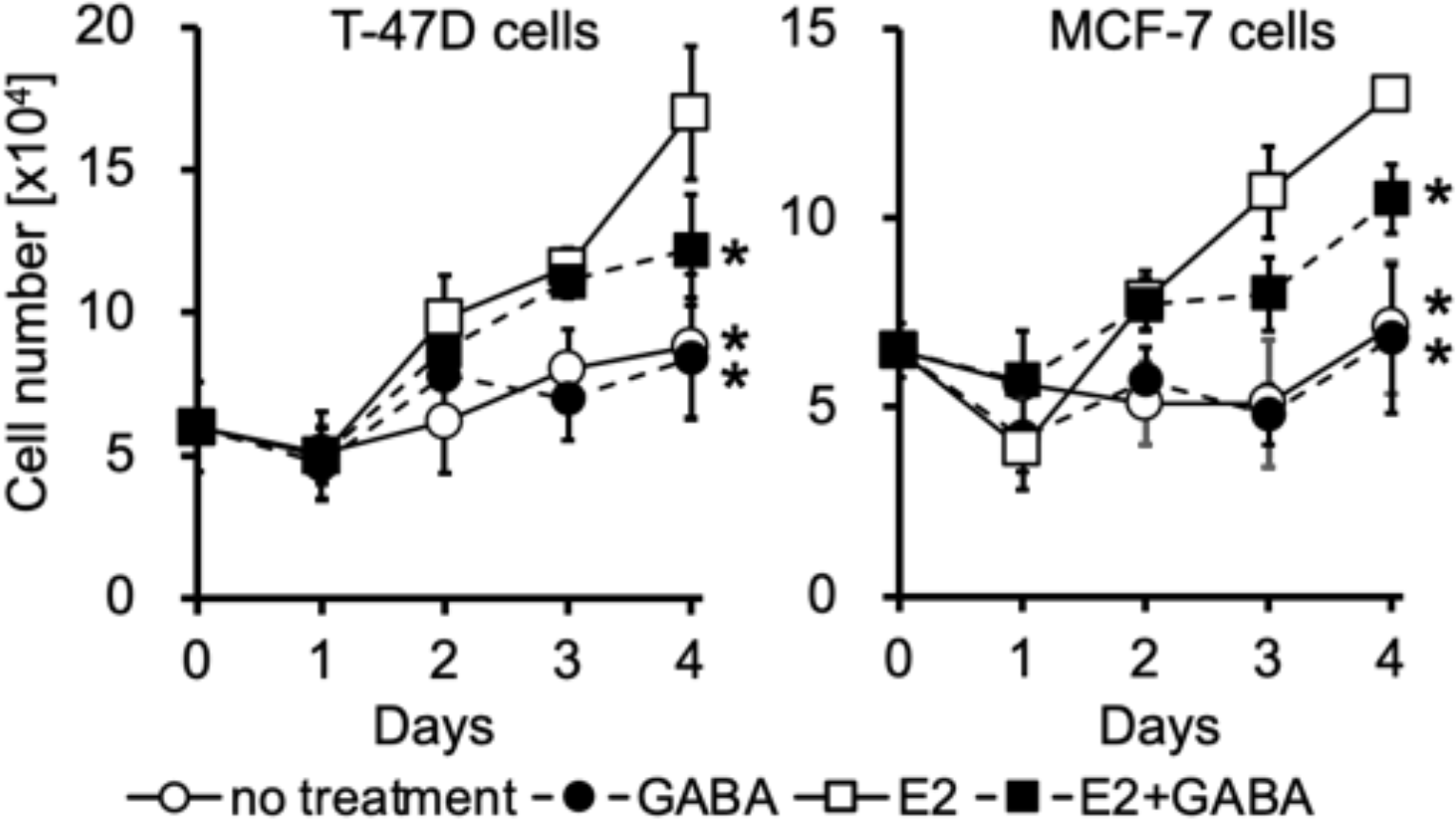
GABA inhibits E2-stimulated cell growth in estrogen receptor positive breast cancer cell lines. The graphs show the results of cell growth assays with T-47D and MCF-7 cells. Asterisks indicate the significant differences to E2 groups at day 4 (*P*<0.05, Dunnett’s test).

E2-stimulation changes the mRNA expression of genes, including protooncogene *MYC* which involves in cell proliferation. We investigated the expression of E2-regulated genes, *TFF1* and *MYC*, with or without GABA (Supplemental Fig. 1). In the results, GABA treatment did not changed in the mRNA levels of the genes. These results suggest that GABA treatment reduces cell growth without changes in gene expression.

### GABA administration reduces mammary dysplasia formation

Daily E2 administration induces mammary dysplasia with loss of biphasic patterns of mammary epithelial cells and myoepithelial cells, abnormal cell growth, and expansion of mammary epithelial cells to the stromal side of a duct in scid mice (23). To investigate the effect of GABA in E2-induced dysplasia formation, we intraperitoneally injected GABA in combination with E2 for 25 days. In the results, we observed dysplasia formation in E2 injected mice, but the majority of mammary ducts in mice injected E2 and GABA exhibited normal morphology (Fig. 4A). We quantified the ratio of dysplasia formation in sections. E2 administration increased ratio of dysplasia formation and GABA reduced it (Fig. 4B).

**Fig. 4.**
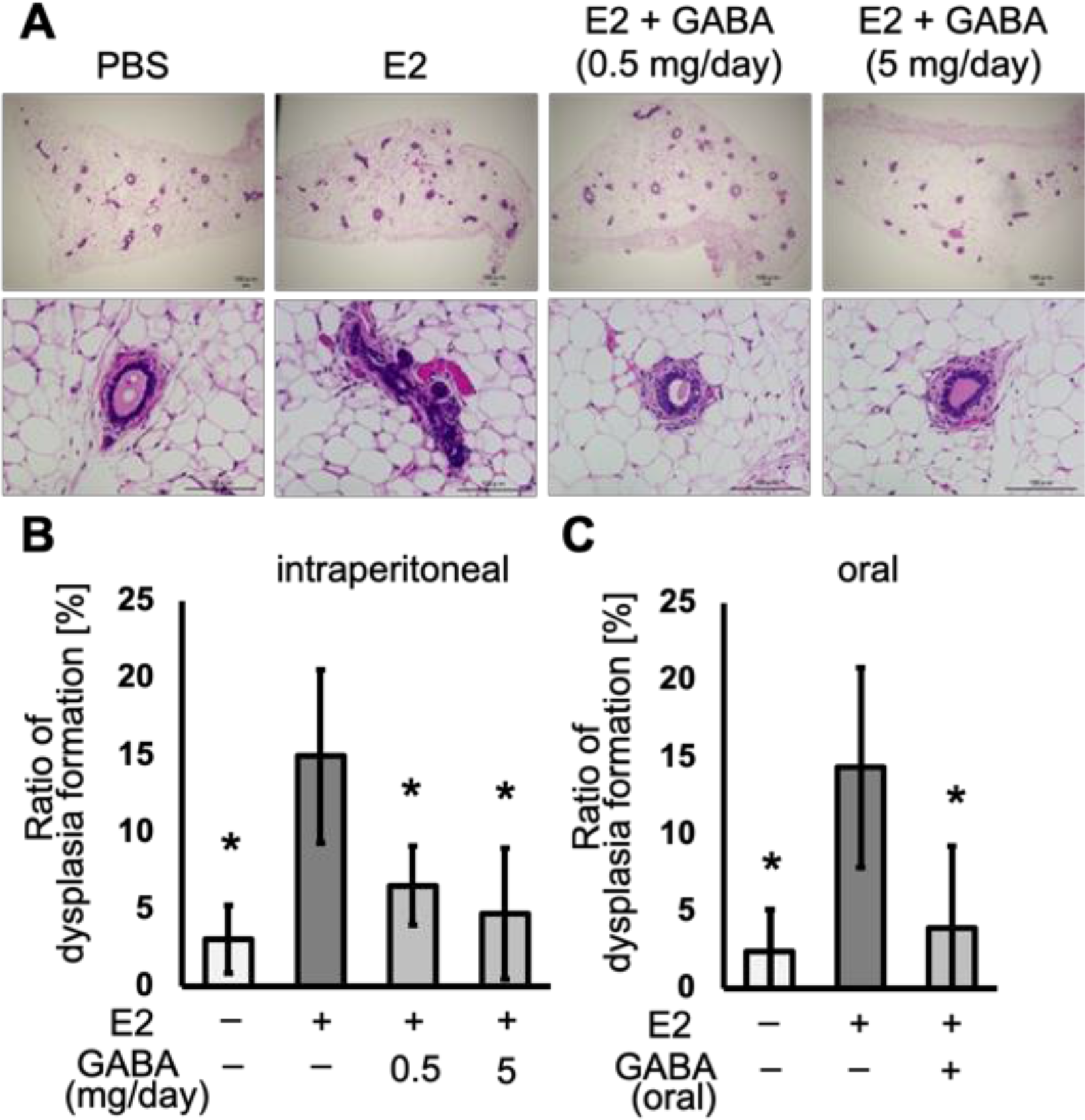
GABA prevents E2-induced mammary dysplasia formation in scid mice. (A) H&E staining of mammary glands are shown. E2 and GABA were intraperitoneally injected. Scale bars indicate 100 μm. (B) Ratios of mammary ducts with dysplasia were quantified in sections of mice injected E2 with or without GABA intraperitoneally. (C) E2 was injected and GABA was orally administered. Asterisks indicate the significant differences to E2 administered groups (*P*<0.05, Dunnett’s test).

To orally administer GABA, we served mice with GABA-containing water. E2 was injected intraperitoneally. The results of histological analyses showed reduction in E2-induced dysplasia formation in GABA orally administered mice (Supplemental Fig. 2) (Fig. 4C). These results indicate that GABA administration reduces E2-induced dysplasia formation in scid mice.

## Discussion

This study showed increase in intracellular GABA level after GABA treatment, although in animal central nervous system, GABA is known as a neurotransmitter which acts as a ligand for a receptor protein without internalization. Our results suggest that GABA is internalized into a cell.

Our cell growth assay showed reduction in E2-stimulated cell growth in GABA-treated groups, suggesting that GABA reduces cell proliferation. Mice early breast cancer induction model system showed reduction in E2-induced dysplasia formation in GABA administered groups. These indicate that GABA prevents early breast tumorigenesis.

Given that estrogen is one of the causes of breast tumorigenesis, selective estrogen receptor modulators (SERM) have a potential to prevent breast cancer. Isoflavones are known materials for breast cancer prevention. The structure of GABA is not similar with SERM and isoflavones, suggesting that the mechanism of breast cancer prevention is not the same between GABA and them. Our findings provides GABA as a novel food material for breast cancer prevention.

GABA exists various foods and easy to obtain from foods. In addition, GABA is added in food products for health. GABA can be stored at room temperature, which facilitate to popularize it widely. On the other hand, breast cancer treatment is constantly advancing, however, due to a large number of breast cancer patients, breast cancer kill a lot of people. Therefore, breast cancer prevention will reduce a number of breast cancer deaths. GABA might contribute to it.

## Methods

### Cell culture

Breast cancer cell lines, MCF-7 and T-47D, were obtained from American Type Culture Collection (Manassas, VA, USA). Cells were maintained with phenol-red-free RPMI 1640 medium (Nacalai Tesque, Kyoto, Japan) supplemented with 10 % charcoal-stripped FBS (Gibco, Waltham, MA, USA) at 37 °C with 5 % CO_2_ in an incubator. For MCF-7 cells, 1 nM of estradiol (E2) (E2758; Sigma-Aldrich, St. Louis, MO, USA) was added in the medium.

### Amino acid analyses

T-47D and MCF-7 cells were cultured with 2 mL of RPMI 1640 medium (Nacalai, Kyoto, Japan) containing 10 % FBS (Sigma-Ardrich, St. Louis, MO, USA) in 35-mm dish (Thermo Fisher Scientific, 130180, NY, USA) for 3 days. After washing with PBS, cells were incubated with medium supplemented with or without 1 μg/mL GABA (Sigma-Ardrich, A5835) for 0.5 h at 37 °C. The media was collected, and the cells were washed with cooled phosphate-buffered saline (PBS), scraped and resuspended with 0.2 mL of cooled PBS. The cells were disrupted by 4 times of freeze-thaw cycles to elute intracellular sample. The sample was centrifuged and the supernatant was collected as an intracellular sample. To measure cellular protein level by Bradford assay for normalization of intracellular GABA level, 0.1 mL PBS was added to the pellet and sonicated by a sonicator (TOMY, UR-21P, Tokyo, Japan) with cooling with ice-cold water.

Samples were separate from proteins with 2.5 % trichloroacetic acid, and filtered through a 0.2 μm membrane filter. Amino acids were detected using the post-column ninhydrin method with an amino acid analyzer (HITACHI, LA8080 SAAYA, Tokyo, Japan) and software OpenLAB CDS Acquisition ver. 2.3 (Agilent Tachnologies, Santa Clara, CA, USA).

### Cell growth assay

For cell growth assay, cells were maintained without E2 for 2 days. T-47D (6 × 10^4^ cells/well) and MCF-7 (6.5 × 10^4^ cells/well) cells were plated in wells of 24-well plate (Techno Plastic Products AG, 92024, Trasadingen, Switzerland) with 0.5 mL of the medium with or without 10 nM E2 and/or 1 μg/mL GABA. To count cells, culture medium was removed, and cells were detached with 0.5 mL of PBS containing 0.5 g/L trypsin and 0.53 mM disodium ethylenediaminetetraacetic acid (EDTA-2Na) at 37 °C. The cell suspension was mixed with an equal volume of 0.4 % trypan blue solution, and cell numbers were measured using an automated cell counter, Countess (Thermo Fisher Scientific).

### Messenger RNA quantification

Total RNA sample was extracted from cultured cells with TRIzol (Thermo Fisher Scientific). RNA was precipitated with 2-propanol and washed with 75 % EtOH. One μg of total RNA was used for cDNA synthesis. Real-time PCR was performed with a reagent FastStart Essential DNA Green Master (Roche Diagnostics, 06 402 712 001, Mannheim, Germany). Relative mRNA level was quantified with ddCt method. The sequences of primers were described previously (23).

### Mouse experiments

Female scid mice were obtained from CLEA Japan, Inc. (Tokyo, Japan) and maintained under specific-pathogen-free condition. Intraperitoneal injection was performed with 27G needle in the morning. Reagents injected were E2 (6 μg/day), and GABA (0.5 or 5 mg/day). For oral administration of GABA, fermented GABA solution (final conc. 1 mg/mL) was used. Fermented solution without GABA was used as a control. Mice were euthanized by cervical dislocation and mammary glands were isolated at 24 h after final injection. The mouse experiments were approved by the Animal Research Committee of Kyoto University, number MedKyo22287. All mice were maintained according to the Guide for the Care and Use of Laboratory Animals (National Institute of Health Publication).

### Histology

Isolated mammary gland was fixed with 10 % formaldehyde neutral buffer solution at 4 °C for overnight, dehydrated and embedded in paraffin. Paraffin-section was cut at 2 μm thick. Deparaffinized section was stained with hematoxylin and eosin (H&E staining).

### Immunostaining

For cultured cells, cells were fixed with 1 % formaldehyde in PBS at room temperature for 15 min. After washing with PBS, permeabilization was performed with 0.1 % Triton-X 100 in PBS at room temperature for 15 min. PBS containing 0.05 % tween-20 (PBS-T) and blocking solution (5 % goat-serum in PBS-T) were used for washing, and blocking and antibody solution, respectively. Primary antibody reaction was performed with anti-GABA antibody (Sigma-Aldrich, A2052, 1/1000 dilution). Secondary antibodies used was anti-rabbit IgG antibody conjugated to Alexa Fluor 555 (Life Technologies Corporation, A31572, Eugene, OR, USA, 1/1000 dilution). The sample was co-stained with phalloidin. Fluoromount-G (SouthernBiotech, 0100-01, Birmingham, AL, USA) was used for mounting.

### Microscopy

H&E staining images were observed with an all-in-one microscope (Keyence, BZ-9000) equipped with objective lenses, ×10 PlanApo (NA: 0.45) and x40 PlanApo (NA:0.95), and collected with software, BZ-II Viewer (Keyence). Fluorescent images were observed with a confocal microscope (Evident, FV10i) equipped with water immersion x60 PlanApo (NA:1.2) objective lens, and collected with FV31S-SW (Evident). ImageJ was used for image processing.

### Statistical analyses

Values were analyzed by Dunnett’s test or Student *t*-test. In *in vitro* analyses, experiments were triplicated. P < 0.05 is considered as statistically significant. Error bars indicate standard deviation.

## Acknowledgement

We thank Dr. Masamichi Kamihira for sharing equipment.

---

**Supplemental Fig. 1.**
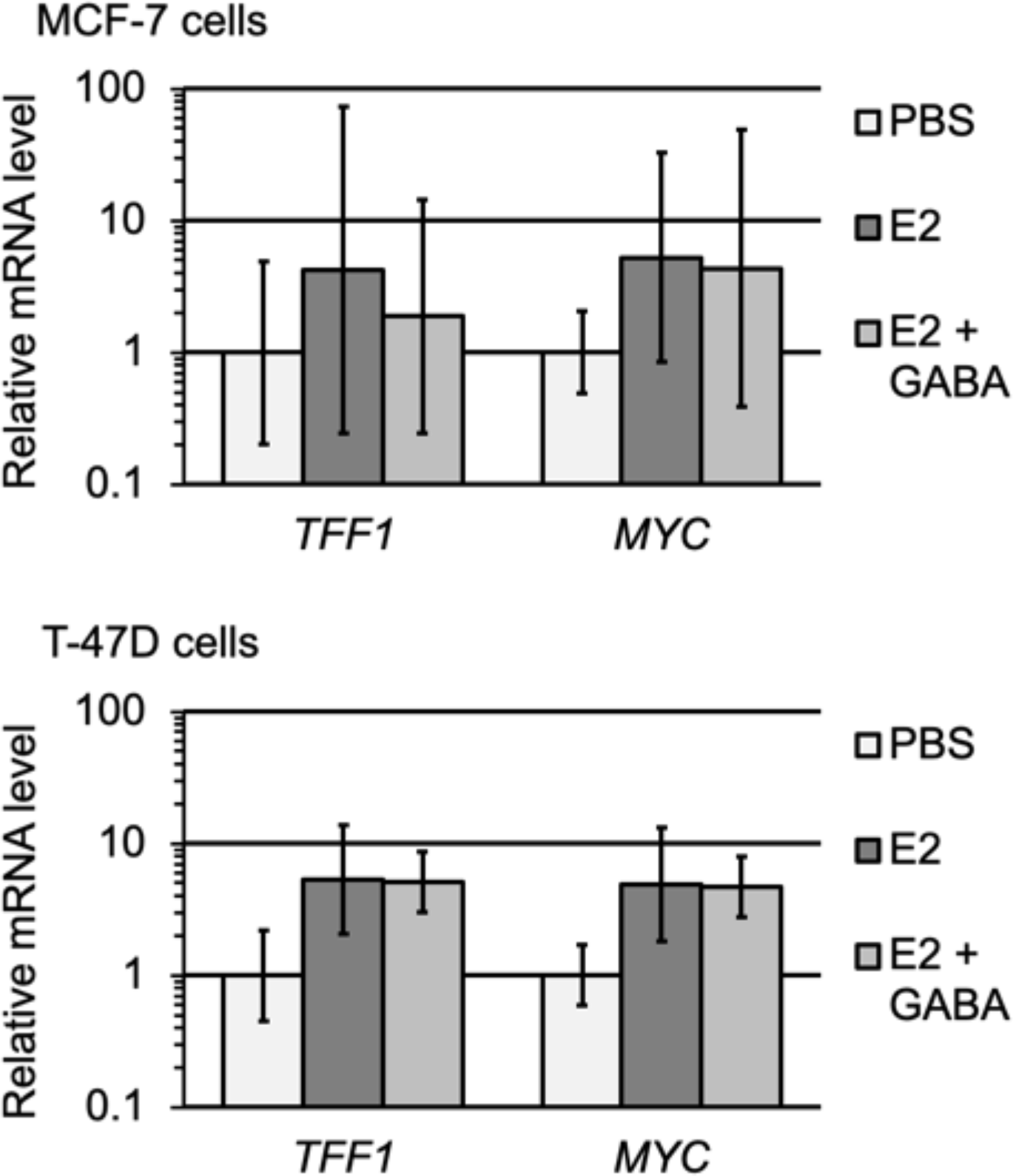
GABA does not affect E2-induced *TFF1* and *MYC* expression. Messenger RNA levels relative to PBS controls were graphed. MCF-7 and T-47D cells were treated with E2 or E2 and GABA.

**Supplemental Fig. 2.**
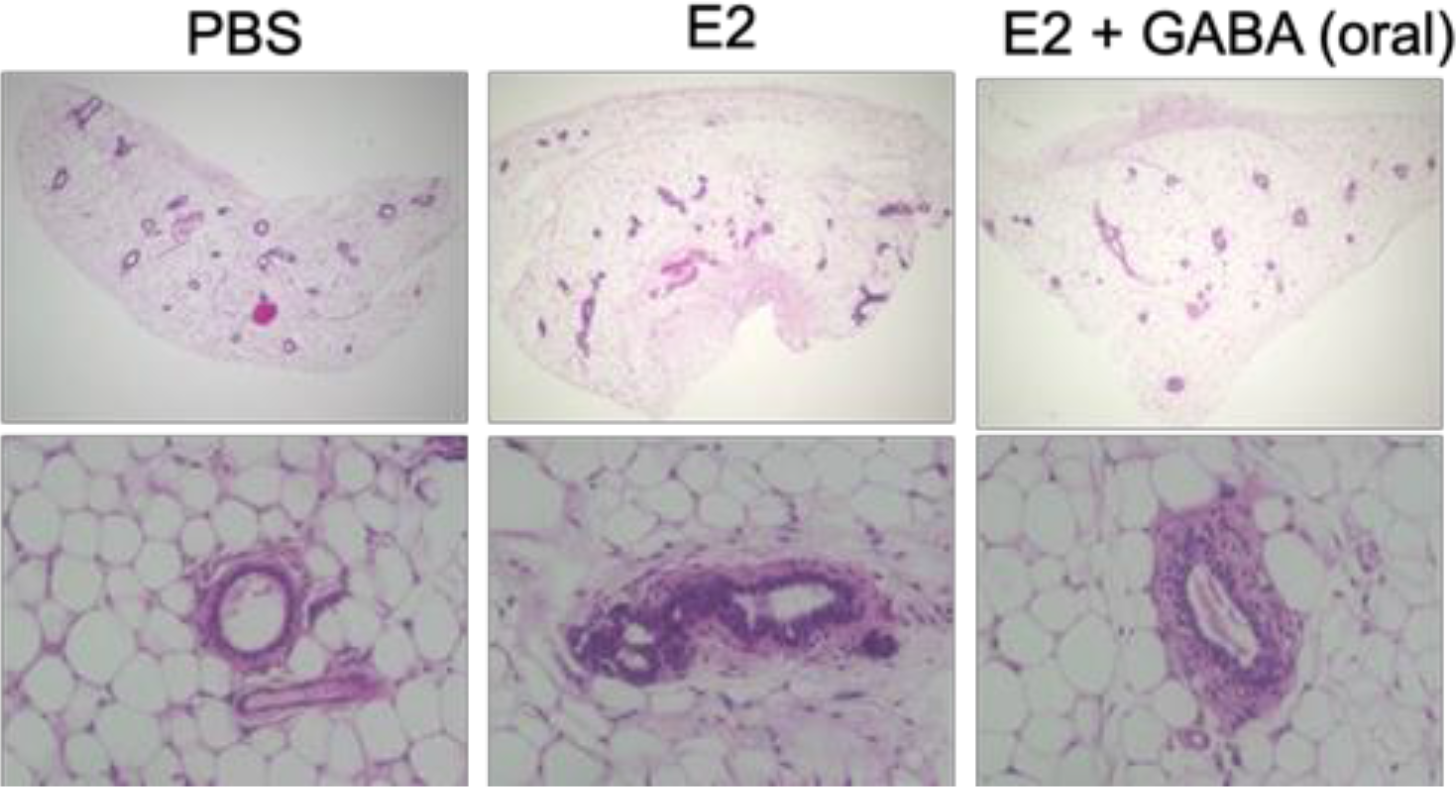
Oral administration of GABA prevents E2-induced mammary dysplasia formation in scid mice. H&E staining of mammary glands are shown. E2 was injected and GABA was orally administered for 25 days.

