## Supplemental Figures for "Gamma-aminobutyric acid treatment reduces estradiol-stimulated breast cancer cell growth and estradiol-induced breast tumorigenesis"

### Supplemental Fig. 1

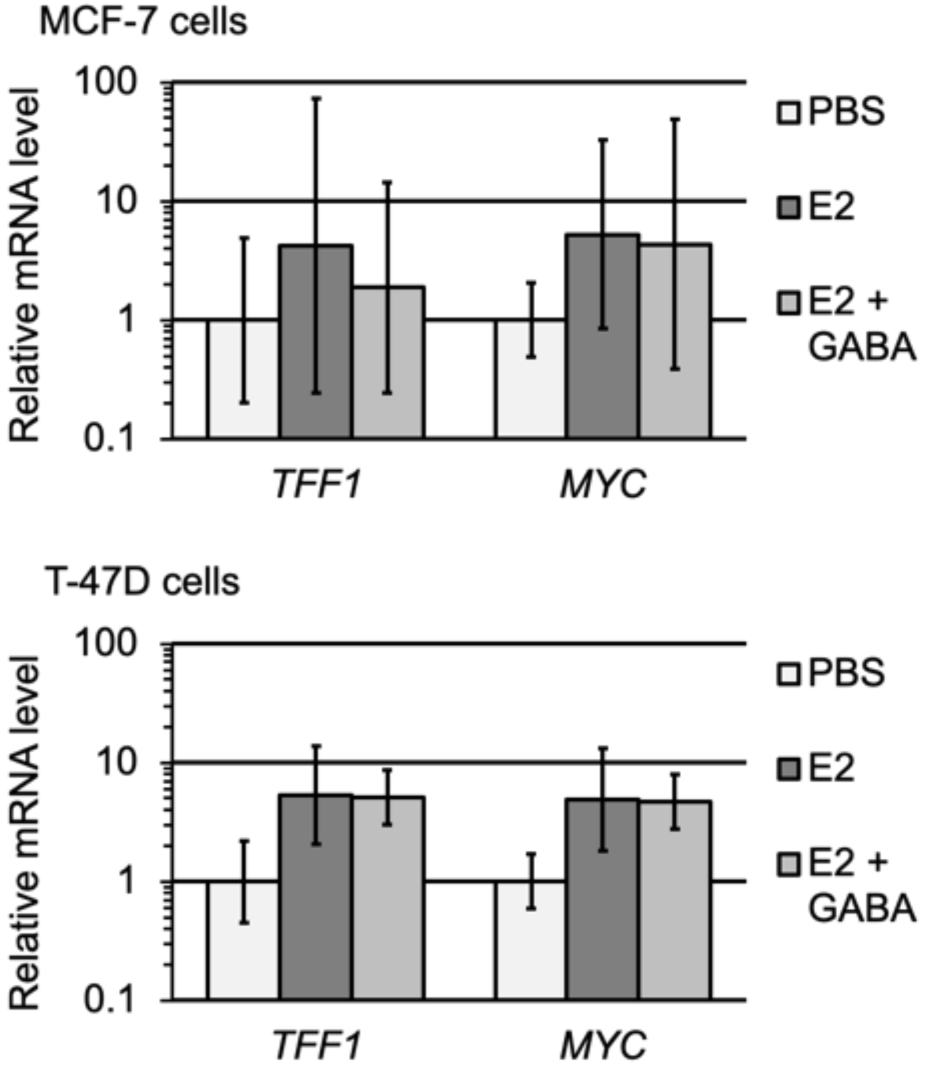


**Supplemental Fig. 1 GABA does not affect E2-induced *TFF1* and *MYC* expression**
Messenger RNA levels relative to PBS controls were graphed. MCF-7 and T-47D cells were treated with E2 or E2 and GABA.

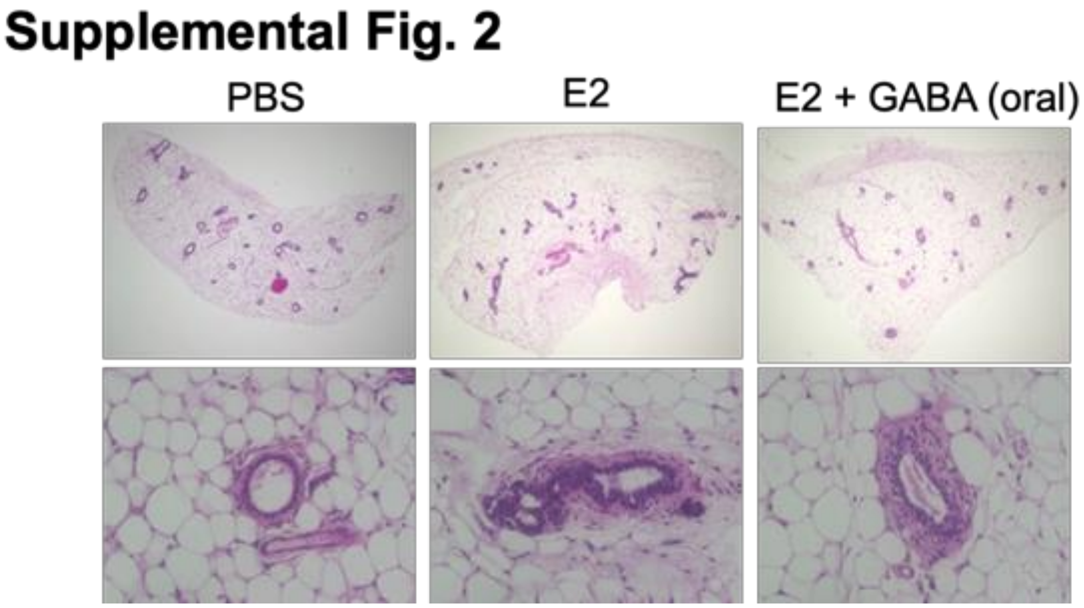


**Supplemental Fig. 2 Oral administration of GABA prevents E2-induced mammary dysplasia formation in scid mice**

H&E staining of mammary glands are shown. E2 was injected and GABA was orally administered for 25 days.
